# Recent evolution of flowering time across multiple European plant species correlates with changes in aridity

**DOI:** 10.1101/2023.04.18.537141

**Authors:** Robert Rauschkolb, Walter Durka, Sandrine Godefroid, Lara Dixon, Oliver Bossdorf, Andreas Ensslin, J.F. Scheepens

## Abstract

Ongoing global warming and increasing drought frequencies impact plant populations and potentially drive rapid evolutionary adaptations. Historical comparisons, where plants grown from seeds collected in the past are compared to plants grown from freshly collected seeds from populations of the same sites, are a powerful method to investigate recent evolutionary changes across many taxa. We used 21-38 year-old seeds of 13 European plant species, stored in seed banks and originating from Mediterranean and temperate regions, together with recently collected seeds from the same sites for a greenhouse experiment to investigate shifts in flowering phenology as a potential result of adaptive evolution to changes in drought intensities over the last decades. We further used single nucleotide polymorphism (SNP) markers to quantify relatedness and levels of genetic variation. We found that, across species, current populations grew faster and advanced their flowering. These shifts were correlated with changes in aridity at the population origins, suggesting that increased drought induced evolution of earlier flowering, whereas decreased drought lead to weak or inverse shifts in flowering phenology. In five out of the 13 species, however, the SNP markers detected strong differences in genetic variation and relatedness between the past and current populations collected, indicating that other evolutionary processes may have contributed to changes in phenotypes. Our results suggest that changes in aridity may have influenced the evolutionary trajectories of many plant species in different regions of Europe, and that flowering phenology may be one of the key traits that is rapidly evolving.

**Highlighted student paper:** We demonstrated that accurately sampled and stored seed collections from conventional seed banks can, with some limitations, be used in a similar way as the increasingly popular resurrection approach sensu stricto. Given the vast availability of seeds stored in seed banks, this opens up a large potential for future research on rapid evolutionary adaptation to changing environmental conditions across a wide variety of taxa suitable for resurrection. Furthermore, this work is unique and novel, as we combine greenhouse experiments with molecular and climatic data to disentangle potential drivers for the observed phenotypic evolution, which, to our knowledge, was never done in resurrection studies.

## Introduction

Over the last decades, climate change has increased dramatically in Europe (IPCC 2021). These changes include both higher temperatures (IPCC 2013) and shifts in precipitation patterns, which often imply increases in the frequency and duration of drought events, as is for instance the case in Southern and Central Europe (Ruosteenoja et al. 2018; Samaniego et al. 2018; Spinoni et al. 2018). Such drought events are often intensified by increased evapotranspiration induced by higher temperatures (Feng and Fu 2013). Plant populations will have to cope with such novel and more stressful environmental conditions (Anderson et al. 2012; Shaw and Etterson 2012; Fleta-Soriano and Munné-Bosch 2016) and are under an increased risk to go locally extinct (Thomas et al. 2004; Urban 2015).

Plant populations may escape local extinction if they respond through adaptive phenotypic plasticity or adaptive evolution (Holt 1990; Hoffmann and Sgrò 2011). Examples for such plastic or evolutionary changes are shifts in phenological events such as leaf-out, flowering onset and time of fruiting, which are key events in plant species’ life cycles and crucial for individual fitness. As phenology is frequently cued by environmental factors (Schwartz et al. 2006; Tang et al. 2016) it is likely that higher temperatures and more frequent droughts influence the timing of phenological events in plant populations. Furthermore, because of the many interactions between plants and animals in ecosystems, shifts in phenology may also impact pollinators, food webs and other ecosystem functions such as productivity or carbon cycling (Reilly et al. 1996, Chmielewski et al. 2004; Cleland et al. 2007; Tang et al. 2016).

Within the last two decades an increasing number of studies showed that plant populations are responding to climate change by shifting their phenology through phenotypic plasticity (Fitter and Fitter 2002; Primack et al. 2004; Panchen et al. 2012) or through adaptive evolution (Franks et al. 2007; Metz et al. 2020). Observational studies using field and/or herbaria data from multiple species illustrated that plant species in general advanced their flowering during the 19th and 20th century and that this is possibly related to increased temperatures. For example, Panchen and colleagues (2012) found that 28 species from the Greater Philadelphia region advanced their flowering by 2.7 days per 1°C rise in monthly minimum temperature within the last 170 years. Although such observational studies across multiple species are crucial to understand the relationship between climate change and shifts in phenology, it remains unclear whether the observed patterns are due to plastic responses or whether plant populations adapted evolutionarily to novel environmental conditions (Franks et al. 2014).

In order to investigate how climate change may have influenced the recent evolution of plant phenology, ancestral plants can be compared with descendant plants when grown in a common garden, using stored seeds and seeds sampled from the same population today (Orsini et al. 2013; Merilä and Hendry 2014; Franks et al. 2018). An increasing number of studies has used this ‘resurrection approach’ to examine rapid evolution in response to increased temperature and drought. There are convincing examples showing that plants advanced their phenology towards earlier flowering in order to avoid drought (Franks et al. 2007; Vigouroux et al. 2011; Nevo et al. 2012; Thomann et al. 2015).

In addition to such adaptive evolutionary processes, genetic differentiation between ancestral and descendant populations can also be influenced by other non-adaptive, i.e. neutral processes such as population bottlenecks, random genetic drift, i.e. the random change of allele frequencies, and immigration of new alleles by gene flow through seed and pollen (Levin and Kerster 1975; Hay and Smith 2003; Ellstrand 2014; Hoban and Schlarbaum 2014). Consequently, both neutral genetic and phenotypic diversity can be affected, potentially leading to changes in genetic variation within populations and genetic differentiation between populations.

Apart from the “Project Baseline”, in which seeds of many species and populations are purposefully collected for future resurrection experiments (Etterson et al. 2016), seeds preserved in conventional seed banks were not sampled from their source populations for such a purpose. However, these seed collections can be used in a similar way and thereby offer untapped resources for environmental change research on recent evolution (Everingham et al. 2021; Rauschkolb et al. 2022a; Rauschkolb et al. 2022b). Moreover, seed banks with stocks from many different species offer a suitable setup for multi-species studies to draw generalised conclusions (*sensu* van Kleunen et al. 2014), potentially revealing parallel evolutionary responses to common drivers.

When using conventional seed bank material for comparisons of populations collected in the past (hereafter referred to as ‘past’) with currently collected populations (hereafter referred to as ‘current’) it is important that the number of stored seeds is sufficiently high - depending on the species, but at least 500 seeds - and that information about sampling locations and sampling protocols is available, and ideally also about the genetic diversity and composition of the seed stocks (Rauschkolb et al. 2022a). However, compared to the current sampling, the sampling strategy of the past is often incomplete, e.g. information on the number of sampled maternal plants, of the average numbers of seeds collected per maternal plant, or how similar these numbers were across maternal plants may be lacking. All of these factors influence the sampled genetic variation and the relatedness of bulked seeds. Molecular markers can be used to gain post-hoc insight in the sampling strategy of the past population and to identify potential non-adaptive genetic effects, like population bottlenecks or random genetic drift, that may have contributed to genetic changes.

In this study we adopted the resurrection approach in a multi-species experiment with seeds from conventional seed banks. We used 21 to 38 years old accessions from 13 different fast-flowering plant species (’past populations’), from Mediterranean and temperate regions of Europe, and compared them with accessions collected at the same sites in 2018 (’current populations’). The studied populations were thus located in two different European bioclimatic systems, with different degrees of recent climatic change. The temperate populations were exposed to stronger recent increases of drought as higher temperatures were coupled with decreased precipitation during the last decades, whereas the Mediterranean populations only experienced increased temperatures with little changes in precipitation. This provided us with opportunities for studying evolutionary trajectories caused by differences in climate change history during the last decades. To investigate potential evolution of phenology in these populations we observed flowering onset and an early growth trait of 15 plants per temporal origin (i.e. past versus current populations) in a greenhouse experiment. To relate changes in phenotypes to changes in aridity, we characterised all populations using the “De Martonne aridity index” (IDM) based on the last six years before seeds of each population were collected in our analyses. In addition, we used genomic single nucleotide polymorphism (SNP) marker data to assess whether sampling strategies between the two different collection times are comparable and whether non-adaptive processes may have contributed to changes of genetic variation between the past and the current populations.

We hypothesised that current populations generally evolved an advanced flowering onset and a shortened life cycle in comparison to the past populations in order to escape from increasing temperatures and drought induced by climate change during the last decades. As the temperate populations experienced stronger relative changes in drought condition during this time, we expected larger evolutionary shifts for these species. We further hypothesised that plants which experienced drier climates within the last six years before seed collection should show stronger advances in flowering onset. Finally, we discuss the potentials and limitations of seed bank collections for resurrection experiments

## Materials and methods

### Seed collection

We obtained seeds collected in the past from the seed banks at the Conservatoire Botanique National Méditerranéen de Porquerolles (CBNMed, Hyères, France), the Meise Botanic Garden (Belgium), the Botanical Garden of the University of Osnabrück (Germany) and the Berlin Botanic Garden and Botanical Museum (Germany). In each case, the seeds of a particular species from a seed bank represented only a single population of origin from the region where the seed bank was located. The only exception was the seed bank from Meise where the species we worked with originated from different regions, Namur and Liège. To be able to study flowering phenology within a two-year greenhouse experiment we chose species which flowered already within the first or second year of cultivation. Furthermore, we chose accessions which were stored for 20 years or longer to allow sufficient time for potential evolutionary processes. Finally, we restricted ourselves to accessions that had sufficient amounts of seeds in the seed banks, for which precise information about the collecting location was available, and which were relatively isolated (e.g. locally rare species growing in small nature reserves) to reduce the chance that the sampled populations were strongly influenced by gene flow and to increase the possibility that the current populations are descendants of the past populations. All seeds from the past were sampled between 1980 and 1997 (Table S1). After bulking the seeds from different individuals, the seeds were cleaned and then stored under appropriate environmental conditions (Table S2) to preserve viability until we received the seed materials in November 2018. To obtain the seeds of the current populations, we collected seeds of all species from the exact same individual sites per species in the spring (Mediterranean species) or summer (temperate species) of 2018. Depending on the available number of fruiting plants, we sampled between 10 and 103 individuals (Table S1) per population and then bulked all seeds, as in the past seed collections.

During the last decades all study sites underwent significant climate change. The Mediterranean species originating from the region of Hyères, southern France, experienced a slight increase in drought in March-July caused by an increase of temperatures by 1.6 °C, although the mean precipitation in these months slightly increased by around 5.5 mm for the period 1991-2020 compared to the baseline period (1961-1990). During the same period average temperatures in Belgium have increased by 0.9 °C in the region of Namur, by 1.3 °C in the region of Liège, by 1.4°C close to Osnabrück and by 1.3°C close to Berlin. In the province Namur mean precipitation in March-July has decreased by about 80 mm, in the province Liège and in Osnabrück by about 35 mm and in Berlin by 2.5 mm (data from the Climatic Research Unit; Camarillo-Naranjo et al. 2019, Harris et al. 2020).

### Germination and experimental design

To reflect species’ natural growing conditions and life cycle, we germinated seeds of 12 Mediterranean species in December 2018 and of 16 temperate species in March 2019. Due to insufficient germination, the number of species that could be successfully transplanted to pots reduced to 9 from the Mediterranean region and to 13 from the temperate region (Table S1). With these 22 species we performed a common garden experiment to compare phenotypic traits between ancestral and descendant populations. After running the experiment for about 1.5 years 13 out of 22 species – 6 from the Mediterranean region and 7 from the temperate region – had flowered (Table 1). Only the data of these species are analysed here.

**Table 1.** Study species used in the experiment with details on the seed bank and the region of origin, year of collection, the 6-year-De Martonne aridity index (mm/°C), the year of flowering onset within our experiment and the number of individuals in the experiment and the flowering rate for past (P) and current (C) population.

| Species | Seed bank/Region | Collection years<br>P / C | 6- year -<br>de<br>Martonne<br>aridity<br>index<br>(mm/°C)<br>P / C | Year of<br>first<br>flowering | Number of individuals<br>and flowering rate<br>P / C |
| --- | --- | --- | --- | --- | --- |
| **Mediterranean species** |  |  |  |  |  |
| *Anthemis maritima* | CBNMed/Hyères | 1992 / 2018 | 25.3 / 35.3 | second year | 15 (93%) / 15 (87%) |
| *Elytrigia juncea* | CBNMed/Hyères | 1994 / 2018 | 26.9 / 35.3 | second year | 14 (64%) / 17 (35%) |
| *Matthiola tricuspidata* | CBNMed/Hyères | 1994 / 2018 | 26.9 / 35.3 | first year | 20 (90%) / 19 (100%) |
| *Medicago marina* | CBNMed/Hyères | 1992 / 2018 | 25.3 / 35.3 | second year | 13 (54%) / 13 (62%) |
| *Plantago crassifolia* | CBNMed/Hyères | 1994 / 2018 | 26.9 / 35.3 | first year | 16 (81%) / 16 (100%) |
| *Plantago subulata* | CBNMed/Hyères | 1997 / 2018 | 32.2 / 35.3 | first year | 13 (85%) / 14 (93%) |
| **Temperate species** |  |  |  |  |  |
| *Centaureum erythraea* | Meise/Namur | 1992/2018 | 39.9 / 32.1 | second year | 14 (57%) / 14 (57%) |
| *Clinopodium vulgare* | Meise/Namur | 1992/2018 | 39.9 / 32.1 | first year | 14 (100%) / 15 (100%) |
| *Dianthus carthusianorum* | Osnabrück/Osnabrück | 1993/2018 | 36.8 / 32.9 | second year | 13 (46%) / 11(82%) |
| *Globularia bisnagarica* | Meise/Namur | 1992/2018 | 39.9 / 32.1 | second year | 12 (75%) / 15 (80%) |
| *Leontodon hispidus* | Meise/Liège | 1995/2018 | 40.2 / 39.2 | first year | 15 (100%) / 15 (100%) |
| *Sanguisorba minor* | Meise/Namur | 1992/2018 | 39.9 / 32.1 | second year | 8 (88%) / 10 (80%) |
| *Silene chlorantha* | Berlin/Berlin | 1980/2018 | 29.8 / 26.5 | first year | 13 (69%) / 13 (54%) |

For germination of the Mediterranean species we placed 100 seeds per temporal origin on 1% water agar in 90 mm Petri dishes, stratified them for one week at 5°C in the dark and then germinated them in a walk-in growth chamber (light intensity = 230 µmol·m^-2^ s^-1^, 50% relative humidity) with a light/dark cycle of 8/16 hours and temperatures of 23/18°C in December 2018. For germination of the temperate species, we placed at least 50 seeds (exact number of seeds Table S1) per temporal origin on moist potting soil (Einheitserde®, BioLine, Pikiersubstrat) in trays (TEKU®, TK1520 18.5 × 14 × 5.1 cm) and stratified the seeds for two months (January and February 2019) at 5°C in the dark. This method reduced fungal infections compared to two months stratification on agar. In March 2019, we transferred the trays with temperate species to the greenhouse and allowed the seeds to germinate at approx. 20 °C under a natural spring daylight regime. After a sufficient number (>25) of seeds per temporal origin (past and current) from Mediterranean and temperate species had germinated we transplanted up to 20 seedlings each into 9 × 9 cm pots (one plant per pot) with a 3:1 mixture of potting soil (Einheitserde®, BioLine, Topfsubstrat Öko torffrei) and sand (0-2 mm play sand, WECO GmbH) and placed them in the greenhouse.

Although for resurrection studies it is recommended to cultivate a ‘refresher generation’ (Franks et al. 2018) to reduce storage and/or maternal effects, we decided to conduct our experiment without a ‘refresher generation’. The reason for this was that we had to expect that some of our study species would not flower within one year and thus the cultivation of a ‘refresher generation’ would have demanded too much time. However, to reduce the impact of storage and maternal effects, we have chosen to correct for differences in germination and early growth by always transplanting pairs of past and current seedlings that were approximately of equal size.

During the whole experiment we blocked all pots of a single species but randomised pot positions every second week within each block. The greenhouse was set to a light/dark cycle of 12/12 hours and temperatures of 20/15 °C as upper/lower limits. Throughout the experiment, we watered the plants sufficiently and regularly (two to three times a week), always giving the same amount of water per species (100-200 mL, depending on the size of a species).

Three to four weeks after transplanting to pots, we measured on each plant one size trait (initial size: shoot length or rosette diameter), depending on the species’ growth form. In April 2019 the first Mediterranean and in May 2019 the first temperate species started to flower. We recorded the day of the year (FT_Ind_) of flowering onset per individual until the end of August 2019. After the first year of our multi-species experiment seven out of 22 species had flowered. For these species, we finished the experiment and scored them as flowered in the first year (Table 1). To stimulate vernalisation we kept the 15 remaining species in a non heated greenhouse from September to November 2019 and let temperatures drop to a minimum of 5°C during December 2019 and January 2020. In February 2020, we cut off dead plant material and transferred the pots back to the experimental greenhouse, repeated the initial size measurements 3 weeks later and recorded the date of flowering per plant until July 2020.

### Statistical analyses

We standardised the flowering records per species and individual by using the following formula:

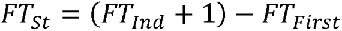

where FT_Ind_ is the day of the year a plant flowered and FT_First_ is the day of the year the first plant of this species flowered.

We analysed our data in two different ways: multiple single-species models and a single multi-species model. For the single-species analyses of flowering time, we used linear models which included temporal origin (past vs. current) as explanatory variable and the initial size, depending on the year of flowering, as covariate. For the multi-species analysis of flowering time, we applied linear mixed-effects models using random slopes of species within the temporal origins as random factor, temporal origin, flowering year (flowered in the first year vs. flowered in the second year) and region (Mediterranean vs. temperate) as fixed factors and the initial size, which we standardised per species to a mean of 0 and a standard deviation of 1, as covariate. We included all interactions of the fixed factors as well as the initial size × flowering year interaction within our multi-species model. In order to improve model residuals, we square-root-transformed the FT_St_ values.

To test whether the temporal origins differed in their early growth, we ran linear single-species models with the species-specific initial size trait (shoot length or rosette diameter measurements) as response variable and the temporal origin as explanatory variable, and a multi-species linear mixed-effects model with the standardised size measurements as response variable and the same fixed and random factors as in the multi-species model for flowering time.

To investigate whether changes in climate during the last decades may have influenced evolutionary shifts in flowering onset in our tested populations we used interpolated monthly mean precipitation and temperature data from the locations of origin obtained from the Climatic Research Unit (Camarillo-Naranjo et al. 2019, Harris et al. 2020). With these data we calculated the “De Martonne aridity index” (IDM, Pellicone et al. 2019) for the last six years before seeds of the population were collected (6-year-IDM). This six year period reduces the influence of environmental fluctuations within the long-term trends while simultaneously covering three to six generations of our experimental species since some of our species flowered within the first, other species in the second year in the greenhouse experiment. For the calculation we used the following formula

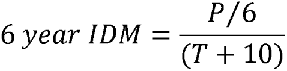

where P is the sum of precipitation (mm) within the 6 years before seed collection and T (°C) is the mean annual temperature over the 6 years before seed collection to which 10 is added to avoid negative values. Thus, lower values of the IDM indicate drier environmental conditions. This index combines changes in temperatures and precipitation, which is important because these factors interact and should be considered together to understand the impacts of climate change on shifts in phenology (Llorens and Peñuelas 2005; Bloor et al. 2010). To test whether the aridity within the last 6 years before seed sampling was related to the mean of flowering time, we ran linear models with the mean of the FT per population as response variable, and IDM, temporal origin, flowering year, region and all interactions of the last three as fixed factors. To test whether the differences in aridity between the two temporal origins are related to the differences in flowering onset we calculated the IDM difference (IDM_Diff_ = IDM_Current_ – IDM_Past_) and the FT difference (FT_Diff_ = FT_Current_– FT_Past_) for each species. We ran linear models with the FT_Diff_ as response variable and the IDM_Diff_, the flowering year, the region and the interaction of the last two as explanatory variables. For all models, we visually checked the residuals for normality and heteroscedasticity. All analyses were done in R (Version 4.0.2) using the packages *plyr* for data management (Wickham 2011) and *lme4* (Bates et al. 2015) and *lmerTest* (Kuznetsova et al. 2017) for running our models using the *lmer()* function and Type III anovas for significance testing.

### Genetic marker analysis

In order to assess whether non-adaptive processes contributed to changes of genetic variation between the past and the current populations, we quantified genetic relatedness, genetic diversity within, and genetic differentiation between the two temporal origins. We used leaf samples of 4-10 (average 9.7) common garden plants per population. We produced genomic single nucleotide polymorphism (SNP) marker data following a ddRAD protocol (Peterson et al. 2012), performed SNP genotyping with dDocent 2.6.0 (Puritz et al. 2014) and conducted SNP filtering following O’Leary et al. (2018). The final data set consisted of 230 samples, between 11 and 20 (average 17.7) per species and between 3 and 10 (average 8.9) per temporal origin, genotyped at between 391 and 2677 (average 1223) bi-allelic SNP loci. For each species, we assessed pairwise genomic relatedness among samples within the past and the current populations using the kinship estimator *r^ß^* (Goudet et al. 2018). The kinship estimator takes values of zero for randomly related pairs, negative values for less closely and positive for more closely related pairs of individuals. We tested for significant differences of relatedness between the two temporal origins by using Wilcoxon rank tests. In addition, we assessed genomic diversity within both past and current populations as allelic richness, Ar, thus correcting for differences in sample size (El Mousadik and Petit 1996). For example, reducing the number of maternal plants from which seeds are collected would increase the average relatedness (*r^ß^>0*) and diminish the allelic richness. Finally, as an overall measure of difference of allele frequencies, we quantified neutral genetic differentiation between the two temporal origins as pairwise *F*_ST_ and tested for significance by bootstrapping over loci. For a detailed account of the SNP genotyping and population genomic analyses, see Methods S2.

## Results

The germination rate across the studied species ranged from 25% to 98%. We observed low germination rates (≤ 30%) for both temporal origins of *A. maritima* and *P. crassifolia* and for the past accession of *E. juncea*, *P. subulata* and *D. carthusianorum*. In general, seeds of the current populations had higher germination rates, but large differences with the past populations (>25 percentage points) were only present in three species (*M. marina*, *P. subulata* and *L. hispidus*, see Table S1).

Plants of the current populations grew faster within the first weeks of the experiment (initial size; *F_1,13.7_* = 4.5, *P* = 0.05; for detailed model outputs of the cross-species analyses here and elsewhere, see Table S3) and this effect was even stronger in the species that flowered in the second year of the experiment (origin × flowering year; *F_1,13.7_* = 10.1, *P* = 0.006).

Regarding flowering onset, species from the same region started to flower evenly distributed during the two growing seasons indicating no clustering by region. Comparing the two temporal origins, we found a significant advance in flowering onset across 13 species, with current populations flowering on average 8.5 days earlier than past populations (*F_1,13.8_* = 5.6, *P* = 0.03). Furthermore, species which flowered in the second year showed a stronger advancing of flowering onset in comparison to the species which flowered in the first year (*F_1,12.8_* = 9.8, *P* = 0.008).

There were substantial differences among species in their magnitudes of evolutionary changes. Only two species showed significant differences between the temporal origins in early growth: *P. subulata* following and *A. maritima* not following the cross-species pattern (Table 2). Furthermore, the individuals of the current populations of five species flowered significantly earlier, whereas seven species showed no significant difference between the two temporal origins and only in *A. maritima* individuals of the past population flowered earlier (Table 2). A generalised linear mixed model (GLMM) with a binomial distribution (Flowering plant: yes vs. no), temporal origin as fixed and species as random factor showed no significant effect of the two temporal origins on the proportion of flowering individuals across species (Chi^2^_1_=1.6, *P* = 0.208).

**Table 2.** F- and P- values of the linear models for the single species analyses showing the effect of the temporal origin on the initial size as well as on the flowering onset, and the effect of the initial size on the flowering onset. The last column presents the shifts in flowering onset in days between the current and the past population.

| Species | Temporal origin (d.F.=1) |  |  |  | Initial size (d.F.=1) |  | Shift in flowering onset (days)<br>(Current - Past) |
| --- | --- | --- | --- | --- | --- | --- | --- |
|  | Initial size |  | Flowering onset |  | Flowering onset |  |  |
|  | *F* | *P* | *F* | *P* | *F* | *P* |  |
| Mediterranean species |  |  |  |  |  |  |  |
| *Anthemis maritima* | 21.3 | <0.001 | 5.1 | 0.03 | 2.5 | 0.19 | 8.3 |
| *Elytrigia juncea* | 0.4 | 0.51 | 0.1 | 0.80 | 0.1 | 0.72 | 1.9 |
| *Matthiola tricuspidata* | 3.3 | 0.08 | 2.2 | 0.15 | 9.4 | 0.004 | -16.8 |
| *Medicago marina* | 0.0 | 0.85 | 0.3 | 0.57 | 0.2 | 0.64 | -2.8 |
| *Plantago crassifolia* | 23.5 | <0.001 | 1.0 | 0.32 | 0.4 | 0.52 | 1.7 |
| *Plantago subulata* | 0.4 | 0.53 | 11.8 | 0.002 | 1.3 | 0.27 | -22.9 |
| Temperate species |  |  |  |  |  |  |  |
| *Centaureum erythraea* | 0.4 | 0.55 | 6.6 | 0.02 | 1.7 | 0.21 | -23.4 |
| *Clinopodium vulgare* | 3.8 | 0.06 | 33.4 | <0.001 | 31.3 | <0.001 | -15.8 |
| *Dianthus carthusianorum* | 0.0 | 0.92 | 0.1 | 0.74 | 0.7 | 0.41 | -2.8 |
| *Globularia bisnagarica* | 0.0 | 0.86 | 4.8 | 0.04 | 0.1 | 0.79 | -6.2 |
| *Leontodon hispidus* | 1.2 | 0.29 | 1.0 | 0.33 | 0.1 | 0.77 | 3.6 |
| *Sanguisorba minor* | 0.5 | 0.51 | 32.9 | <0.001 | 24.8 | <0.001 | -24.7 |
| *Silene chlorantha* | 1.1 | 0.31 | 3.1 | 0.10 | 7.2 | 0.02 | -16.8 |

We also tested whether the initial size of the individuals influenced their flowering onset. Here, we found no relationship across species but we found significant results in four species, three of them (*M. tricuspidata*, *C. vulgare* and *S. chlorantha*) showing larger plants having a more advanced flowering onset and one species (*S. minor*) showing the opposite pattern (Table 2). In addition, an interaction between the initial size and the species’ flowering year (*F_1,276.7_* = 9.3, *P* = 0.002) indicated that the initial size had a stronger positive effect on the flowering onset for the species which flowered in the first year.

Temporal origins of populations from localities which experienced drier environmental conditions (lower 6-year-IDM values) within the last six years before the seeds were collected, flowered earlier (Fig. 1, *F_1_ = 7.4*, *P* = 0.02; partial coefficient of determination between the two variables R^2^ = 0.14). We observed an increase in the aridity in the current temperate populations, indicated by negative IDM_Diff_ in Fig. 2, which corresponded with advanced flowering onset within our experiment (Fig. 2, *F_1_* = 10.4, *P* = 0.01; partial coefficient of determination between the two variables R^2^ = 0.51).

**Fig. 1.**
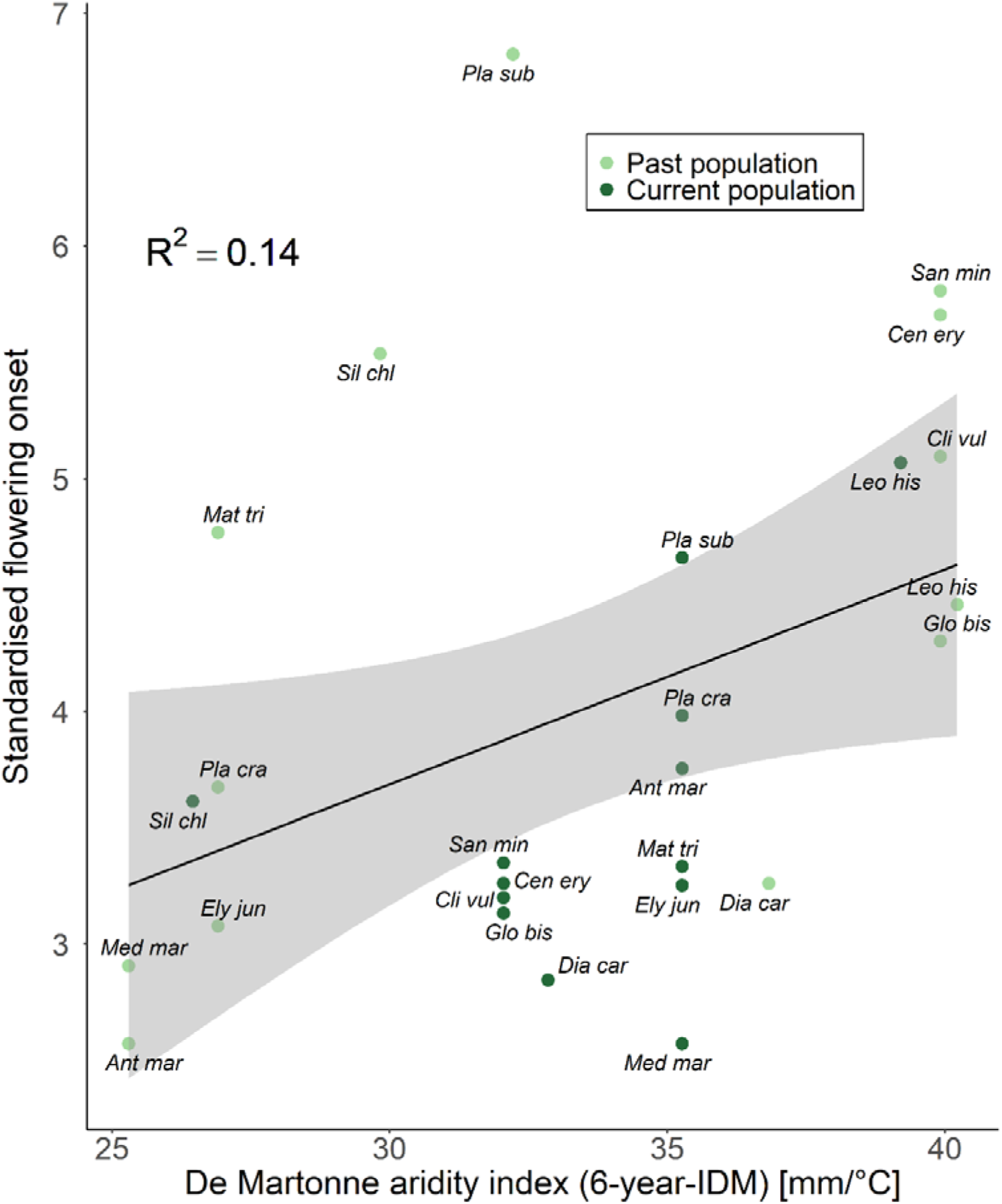
Standardised flowering onset of the 26 different accessions (13 species with past and current population) plotted against the De Martonne aridity index (6-year-IDM). The grey area depicts the 95% confidence interval of the linear regression. R^2^ indicates the partial coefficient of determination between the two variables. *Ant mar* = *Anthemis maritima*, *Cen ery* = *Centaurium erythraea*, *Cli vul* = *Clinopodium vulgare*, *Dia car* = *Dianthus carthusianorum*, *Ely jun* = *Elytrigia juncea*, *Glo bis* = *Globularia bisnagarica*, *Leo his* = *Leontodon hispidus*, *Mat tri* = *Matthiola tricuspidata*, *Med mar* = *Medicago marina*, *Pla cra* = *Plantago crassifolia*, *Pla sub* = *Plantago subulata*, *San min* = *Sanguisorba minor*, *Sil chl* = *Silene chlorantha*

**Fig. 2.**
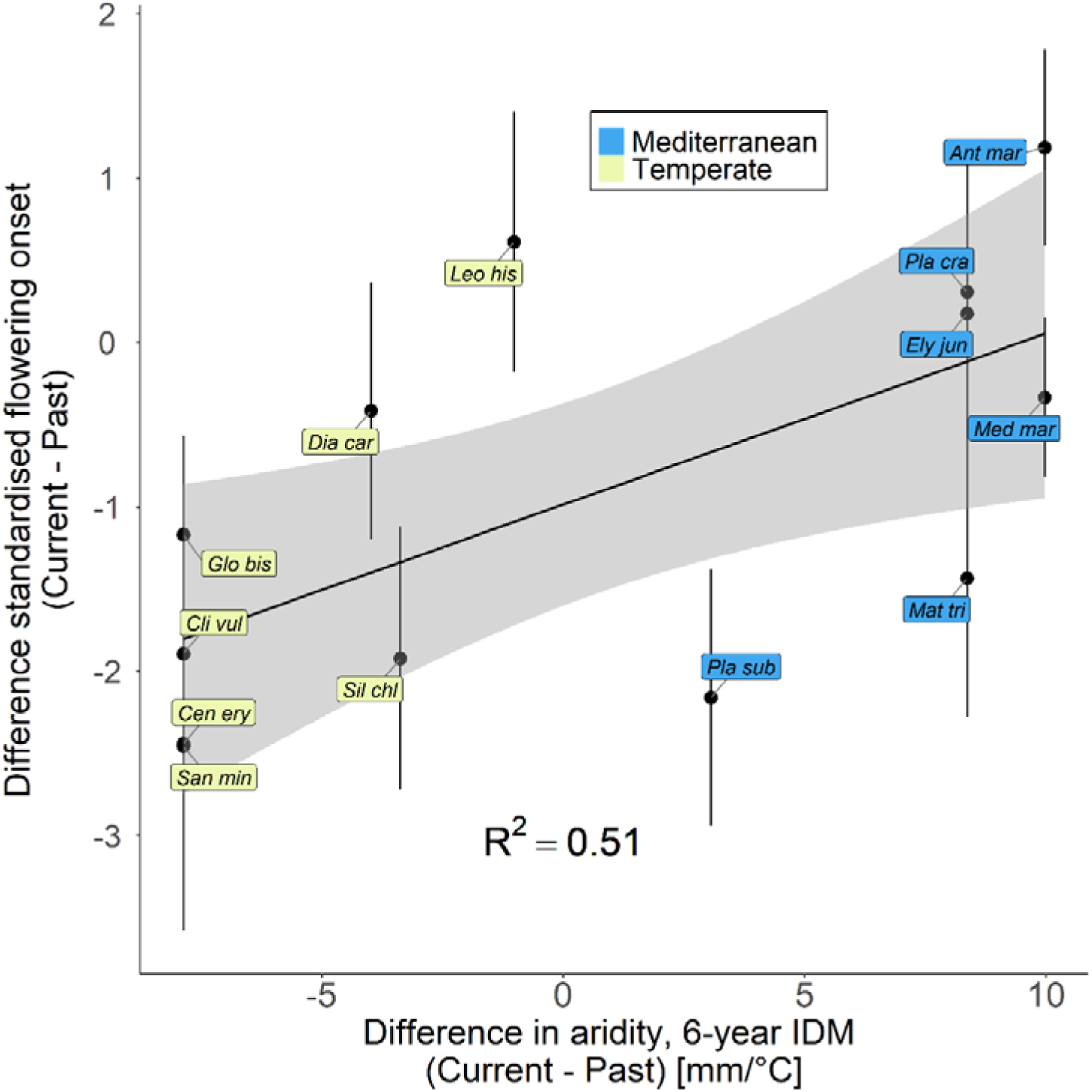
Difference in the flowering onset of the two temporal origins as a function of the difference of aridity (6-year-IDM) between the past and the current population. Negative values therefore mean that the descendant climate is drier. The grey areas depict the 95% confidence intervals of the linear regressions and bars show the sum of the standard errors of both means. R^2^ indicates the partial coefficient of determination between the two variables. For species abbreviations, see caption of Figure 1.

Concerning the relatedness, we found significant differences for the kinship estimator *r^ß^* between current and past populations in six out of 13 species (*M. tricuspidata*, *M. marina*, *P. crassifolia*, *P. subulata*, *S. chlorantha* and *S. minor*, Fig. 3a). The current populations of *P. crassifolia*, *P. subulata*, *S. chlorantha* and *S. minor* had lower *r^ß^* values, whereas the current populations of *M. marina* and *M. tricuspidata* showed higher *r^ß^* values compared to the past populations. In addition, we found large differences in the allelic richness (*Ar*_des._ / *Ar*_anc._ > 1.1 or < 0.9) in four of the aforementioned species (*M. marina, P. subulata, S. minor* and *S. chlorantha;* Fig. 3b) which was often accompanied by large *F*_ST_-values (Table 3). For *M. marina Ar*_des._ / *Ar*_anc._ was < 0.9 indicating higher diversity in the past population, whereas for the remaining three species the current populations showed higher diversity compared to the past populations.

**Fig. 3.**
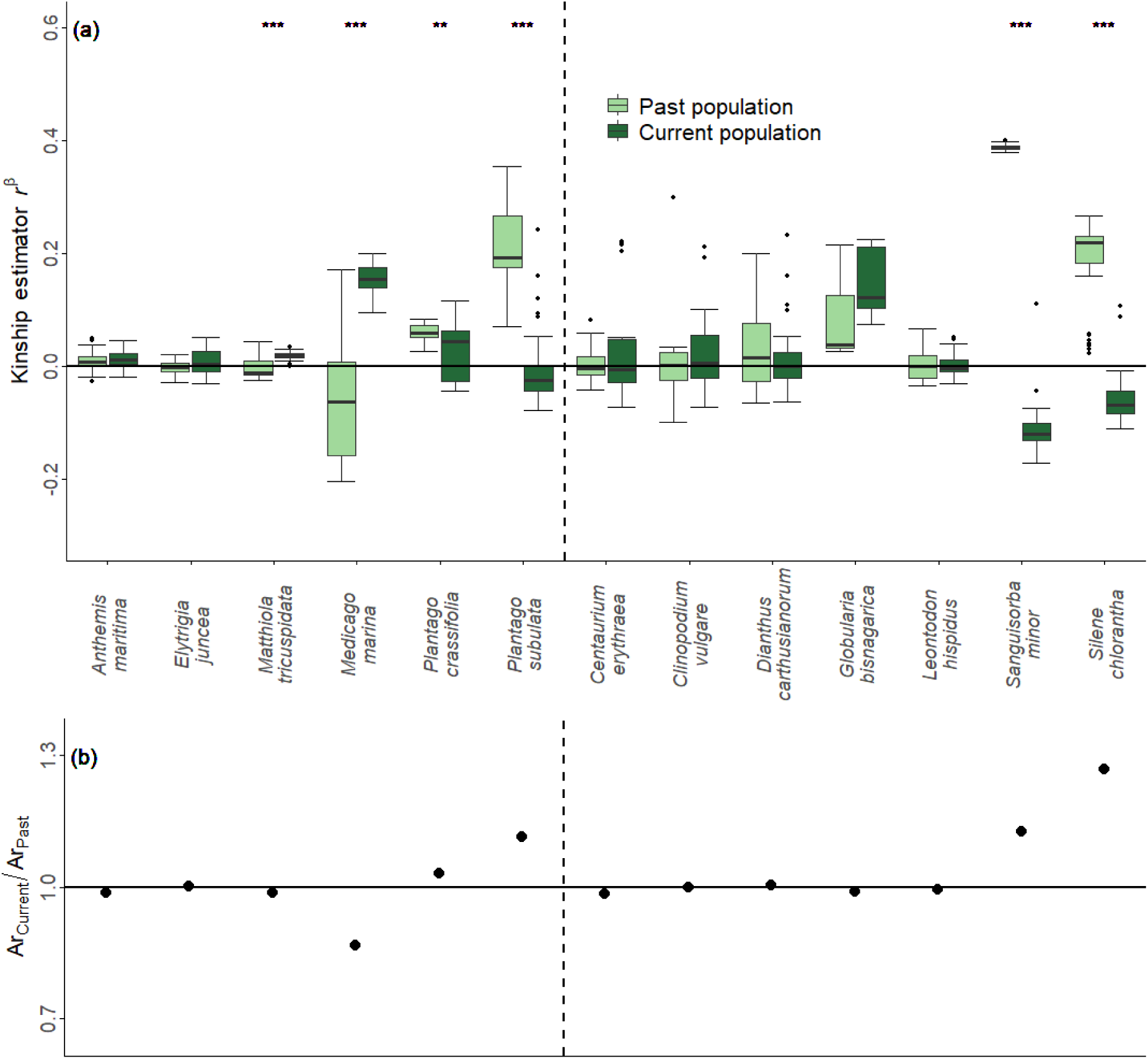
Comparative genomic SNP marker analysis for the past/current population pairs of 13 plant species: six Mediterranean species and seven temperate species separated by the dashed line. (a) kinship estimator *r^ß^*; values of zero indicate random relatedness (*P* < 0.01**, *P* < 0.001***); (b) relative genomic diversity (allelic richness) of the populations (*Ar*_Current_ / *Ar*_Past_); values <1 indicate a reduction of genomic diversity and values > indicate an increase of genomic diversity between past and current populations.

**Table 3.** Overview of molecular data results of the investigated species including information about sample size for the past (P) and the current (C) populations, the number of loci, and genetic differentiation between past and current populations expressed as *F*_ST_, including its significance based on bootstrapping over loci (**, *P* < 0.01).

| Species | Number of samples<br>P / C | Number<br>of loci | $F_{ST}$ significance |
| --- | --- | --- | --- |
| **Mediterranean** |  |  |  |
| *Anthemis maritima* | 10 / 10 | 587 | 0.021** |
| *Elytrigia juncea* | 9 / 10 | 806 | 0.004 |
| *Matthiola tricuspidata* | 10 / 9 | 612 | 0.013** |
| *Medicago marina* | 10 / 10 | 553 | 0.088** |
| *Plantago crassifolia* | 10 / 10 | 1435 | 0.085** |
| *Plantago subulata* | 7 / 10 | 2272 | 0.159** |
| **Temperate** |  |  |  |
| *Centaureum erythraea* | 7 / 7 | 1699 | 0.034** |
| *Clinopodium vulgare* | 6 / 8 | 1047 | 0.040** |
| *Dianthus carthusianorum* | 9 / 8 | 2044 | 0.057** |
| *Globularia bisnagarica* | 3 / 8 | 391 | 0.261** |
| *Leontodon hispidus* | 10 / 10 | 1288 | 0.002** |
| *Sanguisorba minor* | 10 / 9 | 2677 | 0.276** |
| *Silene chlorantha* | 10 / 10 | 485 | 0.114** |

## Discussion

### Phenological differentiations between past and current plants

Our first aim was to investigate whether flowering onset in plant populations from Mediterranean and temperate origins has evolved during the last decades. Indeed, we found that across the 13 studied species, current populations flowered earlier than their 21-38 years older past populations, which was confirmed at the species level for five species and contradicted by only one species (*A. maritima*). This observation is in line with other studies demonstrating that advanced flowering may reflect adaptation to an increasingly drier climate (Franks et al. 2007; Kigel et al. 2011; Metz et al. 2020) since early flowering individuals have a better chance to escape summer droughts. This relationship between drought and advanced flowering is further supported by our comparison of the aridity index of the sampling sites and flowering onset (Fig. 1) which revealed that temporal origins experiencing drier conditions in the years prior to sampling also flowered earlier. This relationship is especially interesting in the Mediterranean species, as the aridity index of the current populations (2012- 2018) was always higher, which mean less dry conditions, than that of the past populations. As a result, this leads to weaker or inverse (*A. maritima*) responses in some of the studied plant populations compared with the general trend that current populations advanced their flowering (origin × region interaction for flowering onset; Fig. 2). For some temporal origins, such as the current population of *E. juncea* or the past population of *D. carthusianorum*, a substantial portion of the living individuals did not flower over the course of the experiment. Here we cannot rule out a potential ‘invisible fraction’ bias (Weis 2018) as these non flowering individuals could have flowered earlier or later than the other individuals of the same temporal origin if we had kept them for subsequent growing seasons.

The advances in the flowering onset for the current populations of some species (*P. subulata*, *C. erythraea* and *S. minor*), measured in the common garden, were very large (>20 days) with a rate of shifting of about 1 day per year, which is remarkably high in comparison to other studies. For example, Metz and colleagues compared populations of *Biscutella didyma* which experienced artificial watering regimes for 10 years and found earlier flowering of 3-4 days (rate: 0.3-0.4 days per year) in the drought-treated population in comparison to the control population when grown in a common garden (Metz et al. 2020). These ranges of flowering shifts were also confirmed in other studies with *Hordeum spontaneum* showing a shift of 0.39 days per year (Nevo et al. 2012) or with *Centaurea cyanus* shifting 0.17 days per year (Thomann et al. 2015). However, Franks and colleagues detected shifts of 1.8 and 8.6 days comparing ancestors of *Brassica rapa* collected in 1997 with their descendants collected in 2004 after several years of drought, translating to rates of 0.27 and 1.22 days per year (Franks et al. 2007), which would be in line with our results for *P. subulata*, *C. erythraea* and *S. minor*.

Cross-species analyses showed that the individuals of the current populations grew faster than those of the past populations within the first three weeks of the experiment, suggesting that current populations have accelerated their life cycle not only through advanced flowering but also through faster growth. To reduce the influence of maternal effects, we always transplanted seedling pairs of similar size from both temporal origins. This procedure may have introduced a bias, as we only selected a subset of the seedlings, i.e. those seedlings which had a similarly sized seedling from the other temporal origin available. By applying this, we may have diminished possible true differences between the past and current populations in early growth and possibly even other traits later in the life cycle. In fact, this conservative bias may explain why we only found a significant difference in initial size between the temporal origins (measured 3-4 weeks later) in two out of 13 species and it may have obscured differences in flowering time.

Herbaceous plants have to reach size thresholds before they start flowering (Mooney et al. 1986; Sun and Frelich 2011) and therefore fast-growing plants can flower earlier. However, in our experiment plants of only three species (*M. tricuspidata*, *C. vulgare*, *S. chlorantha*) with larger initial size in current vs past populations also flowered earlier and there was no cross-species effect. Interestingly all three species flowered in the first year of the experiment and we also found an initial size × flowering year interaction showing that the flowering onset of species flowering in the first year is more strongly influenced by the initial size. We therefore argue that the connection between fast growth and early flowering is more important for the species that flowered in the first year of our experiment as these are comparable to ruderal or annual species which rapidly fulfill their life cycle by fast growth and early flowering (Grime 1977; Fitter and Fitter 2002). In contrast, the species that flowered in the second year are less strongly influenced by the initial size. As those species experienced a period of stasis over winter, their flowering onset might be synchronised by the start of the growing period after winter, reducing the importance of the initial size. To investigate whether the increased growth of the current populations within the first three weeks supported the advanced flowering, we should have measured other traits like size of the plants at flowering or the relative investment into vegetative and reproductive biomass. We also found that the species which flowered in the second year did so within a shorter time after cutting dead plant material after the winter and moving the pots to the warm greenhouse compared to the time between transplanting and flowering of species that flowered in the first year. This may be due to species differences, due the fact that these species could draw on stored metabolites in spring, or due to differences in environmental conditions in the greenhouse between the two years.

### Non-adaptive genetic changes between past and current plants

A fundamental question with our approach is the genealogical relationship between the newly collected seeds and those collected >20 years ago. Apart from selective processes, neutral processes can affect genetic variation and genetic relatedness among plants, which can be revealed when comparing stored seeds from the past and freshly collected seeds. Molecular genetic markers can help to elucidate this question. Population bottlenecks occurring after the seed collection in the past could have led to genetic drift, thus strongly changing the genetic makeup, in particular reducing genetic variation and increasing relatedness (Hay and Smith 2003; Hoban and Schlarbaum 2014). Similarly, gene flow could have led to immigration of new alleles from adjacent populations, thus increasing genetic variation and reducing relatedness (Levin and Kerster 1975; Ellstrand 2014). In the extreme case a population collected >20 years ago could have gone extinct in the meantime and the site could have been recolonised from an external source, thus erasing any ancestor-descendant relationship. Furthermore, the genetic composition of the current populations could have been affected by temporal variation in population size (e.g. smaller size leads to increased genetic drift) and by soil seed bank dynamics through stochastic recruitment. It can also be hypothesised that in self-compatible species the degree of outcrossing and thus the relatedness of seeds may vary among years, e.g. depending on pollinator activity. All these processes, although affecting the genetic pattern, still only involve neutral processes occurring in nature within and among populations. In contrast, artificial selection during sampling may affect the genetic variation encompassed in a seed collection, in particular the number of maternal plants from which seeds were collected, their level of inbreeding and the portion of seeds from each maternal plant in the bulked seedlot (Hay and Smith 2003; Hoban and Schlarbaum 2014).

In six species we found significant difference of *r^ß^* between the two temporal origins, accompanied with strong differences in the allelic richness in four of them. A significant increase in relatedness was found in *M. marina* and *M. tricuspidata*. In contrast to *M. marina* the difference of *r^ß^* between the past and the current population in *M. tricuspidata* was rather small and both temporal origins showed the same allelic richness. We therefore conclude that this difference is a statistical artefact rather than a result of neutral evolutionary processes or artificial selection during sampling. In contrast, for *M. marina* we hypothesise that population bottlenecks and disturbance have occurred between the two sampling dates in this population, thus strongly reducing genetic variation. For *P. subulata*, *S. chlorantha* and *S. minor* a strong and for *P. crassifolia* a slight gain of alleles, accompanied by strong and slight reductions of relatedness were found, respectively. For these populations, gene flow, immigration, metapopulation processes, or different sampling schemes may have led to strong genome wide genetic differentiation between the two generations. For instance, we could expect that less or more closely related plants were collected in the past (Hay and Smith 2003; Hoban and Schlarbaum 2014). With regard to this, we cannot rule out invisible fractions (Weis 2018) for *P. subulata,* as the germination rate of the past population was clearly smaller (29%) in comparison to the current population (78%). Thus, it is quite likely that the investigated individuals only represent a subset of the existing phenotypes. Alternatively, demographic events like gene flow from nearby populations may have caused the increased diversity in the current population (Levin and Kerster 1975; Ellstrand 2014). Summarising, for five species (*M. marina*, *P. crassifolia, P. subulata, S. chlorantha* and *S. minor*) as both neutral processes and selection by aridity due to climate change were active, the observed phenotypic change cannot unequivocally be attributed to adaptive evolution. Thus, for these species it remains unclear whether the observed differences in flowering onset resulted from selective processes, possibly related to escape from summer drought, or were due to other processes that changed the genetic composition between the two time points.

In the rest of the species, i.e. *A. maritima*, *E. juncea*, *C. erythraea*, *C. vulgare*, *D. carthusianorum*, *G. bisnagarica*, *L. hispidus* and, despite of the significant difference in *r^ß^*, *M. tricuspidata*, both past and current populations showed low *F*_ST_ values and similar levels of genetic variation and relatedness suggesting that in these species neutral processes are unlikely to have affected the populations. We can therefore suspect that in these species the current populations are probably related in direct line with the past populations, suggesting that selection is the main and dominant driver of evolution. Having said that, spatial patterns of genetic variation in these species would need to be investigated to make a conclusive assessment. Nevertheless, for these species it is likely that the differences in flowering onset between past and current populations are the result of directional selection imposed by changes in climate (Franks et al. 2007; Kigel et al. 2011; Metz et al. 2020). In fact, in another study, *Q*_ST_-*F*_ST_ comparisons with four species and a similar design as the current study confirmed that significant differences in flowering onset between the two temporal origins were due to past selection processes (Rauschkolb et al. 2022b).

### Using seed bank material for historical comparisons

As we used a common garden approach with controlled conditions for both temporal origins we are confident that the observed differences between past and current populations are due to – potentially adaptive – evolutionary processes and not due to plastic responses. Since flowering phenology is often highly responsive to environmental cues or threshold values, flowering time differences from observational studies may to a large extent reflect plastic responses (Fitter and Fitter 2002; Primack et al. 2004; Panchen et al. 2012). Our historical comparison overcomes this disadvantage but also has some weaknesses. First of all, the material from the seed banks was not collected with the aim to conduct resurrection approach experiments. Therefore, the sampling design and effort may have been different between the past and the current populations and invisible fractions could appear during storage. Also, seeds in the past were bulked, which prompted us to do the same with currently sampled seeds in 2018. Thus, in three species (*P. subulata*, *S. minor*, *S. chlorantha*,) the relatedness strongly decreased over time likely indicating different sampling schemes. For the other species we are confident that the seeds could be used in our historical comparison in a way similar to standard resurrection studies. Reasons for this are (1) the presumed aim of the collectors to maximize the number of sampled individuals in the past, (2) the large amount of seeds within the stored lots and (3) the similar relatedness of the two temporal origins, which was shown by molecular marker results. The latter result provides further support for similar sampling procedures in the past compared to today and indicates that a sufficient amount of seeds was sampled during both periods (Rauschkolb et al. 2022a). Furthermore, in contrast to standard resurrection studies, we did not grow a ‘refresher generation’ before we conducted our main experiment, which would have taken up to 3 years given that a portion of the species only started flowering in the second year. We therefore, in contrast to recommendations for resurrections studies, cannot exclude potential storage or maternal effects (Franks et al. 2018). However, since maternal plants mainly have strong influence on seed characteristics, such as seed provisioning, maternal effects may play an especially important role during germination (e.g. germination time) and seedling establishment (e.g. survival and early growth) (Galloway 2001; Gimeno et al. 2009) and diminish rapidly with plant development (Hereford and Moriuchi 2005; Bischoff and Müller-Schärer 2010). Moreover, we partially accounted for maternal effects by transplanting pairs of past and current seedlings that were approximately of equal size (Latzel 2015). Finally, similar differences in flowering onset between the two temporal origins could be found in another study by Rauschkolb and colleagues (2022b), who conducted an experiment with four species also included in this study. Since they used the same seed material but additionally grew a ‘refresher generation’, the similar results presented here may suggest that the cultivation of a ‘refresher generation’ is not decisive when observing late life stages in resurrection experiments. Thus, we are confident in the validity of our approach using conventional seed bank material, which has the advantage that we could include 13 species with widely different ecologies in this study, allowing us to draw cross-species conclusions on parallel evolution of flowering time.

## Conclusion

It is predicted that ongoing changes in the aridity due to climate change influences the evolution of plant populations. So far, experiments testing this have focused mostly on single species and studies investigating a large set of species are rare. Such a multi-species approach is valuable because it can demonstrate parallel evolutionary responses to broad-scale environmental changes and can reveal different evolutionary trajectories due to different selection pressures in different regions. In this study, we adjusted the ‘resurrection approach’ and used historical comparisons comparing plants grown from seeds collected in the past and their current relatives collected in 2018 of 13 different species from two different biogeographic regions in Europe.

We observed that current populations accelerated their reproductive cycle by faster growth and advanced flowering and that flowering time partly correlated with the short-term climatic backgrounds of the temporal origins. Although we consider for some species that the observed phenotypic differences between the two temporal origins may be due to random changes in genetic composition between the two different time points, we still found consistent results across species, the temporal origins and their climatic developments. Therefore, it would be unlikely that part or all observed patterns in plant responses were due to neutral evolutionary processes (i.e, drift and gene flow) or unintentional selection during seed collection campaigns and experimental set up. Thus, we conclude that selection has driven most of these evolutionary changes. Since we only used one population per species, we admit that our results can say little about the individual species per se. However, taking all species together, our results suggest that climate change may have influenced the evolutionary trajectory of many plant populations in different regions of Europe in parallel over the last few decades. With our study we also demonstrated that historical comparisons using seed bank material are a powerful tool for studying rapid evolution in multiple plant species. We recommend that, if possible, future studies should use seeds from refresher generations to minimise possible maternal or storage effects (Rauschkolb et al. 2022b). Furthermore, fitness measures should be incorporated to disentangle adaptive from non adaptive and maladaptive responses to recent climate change.

## Supplementary Information

The online version contains supplementary material available at …

## Acknowledgements

We thank Nicolai Friesen and Jaroslav Kloster (University of Osnabrück) for their help with the seed collections in the Osnabrück region, Thomas Duerbye and Elke Zippel (Berlin Botanic Garden and Botanical Museum) for their help with seed collections in Berlin, authorities in France, Belgium and Germany for issuing the permits for seed sampling, Lisa Ehmann, Lisa Henres, Christiane Karasch-Wittmann, Denise Klein, Erik Lemke and Eva Schloter for supporting the work in the greenhouse, Madalin Parepa (University of Tübingen) for discussions on data analyses, and Stephan Schreiber and Matthias Bernt (UFZ Halle) for their help with sequencing and bioinformatic analyses.

## Declarations

### Authors’ contributions

RR, AE and JFS conceived the experiment. RR, AE and JFS designed the experiment. RR, SG, and LD conducted fieldwork and RR performed the experiment. RR collected data and performed data analysis with input from OB, AE and JFS. WD performed the molecular analyses. RR wrote the manuscript with input from all co-authors.

### Availability of data and material

The experimental data are available from the digital library of Thüringen (DLT): https://www.db-thueringen.de/receive/dbt_mods_00050983. Raw sequence data have been deposited in the European Nucleotide Archive (ENA) at EMBL under accession number PRJEB47887 (https://www.ebi.ac.uk/ena/browser/view/PRJEB47887) with individual accession numbers ERS7667629 to ERS7668109

### Code availability

Not applicable.

### Ethic approval

Not applicable.

### Consent to participate

Not applicable.

### Consent for publication

Not applicable.

## Footnotes

1 RR, AE and JFS conceived the experiment. RR, AE and JFS designed the experiment. RR, SG, and LD conducted fieldwork and RR performed the experiment. RR collected data and performed data analysis with input from OB, AE and JFS. WD performed the molecular analyses. RR wrote the manuscript with input from all co-authors.

