## Supplementary material for "Recent evolution of flowering time across multiple European plant species correlates with changes in aridity": Table S1, S2, S3, Methods S1

**Electronic supplemental material 1 (ESM1)**

**Parallel evolution of advanced flowering across multiple European plant species in response to increased aridity over the last decades**

Robert Rauschkolb^1,2*^, Walter Durka^3,4^, Sandrine Godefroid^5^, Lara Dixon^6^, Oliver Bossdorf^2^, Andreas Ensslin^7^, J.F. Scheepens^8^

^1^ Institute of Ecology and Evolution with Herbarium Haussknecht and Botanical Garden, Department of Plant Biodiversity, Friedrich Schiller University Jena, Germany, Philosophenweg 16, 07743 Jena, Germany

^2^ Plant Evolutionary Ecology, Institute of Evolution and Ecology, University of Tubingen, Auf der Morgenstelle 5, 72076 Tübingen, Germany

^3^ Department of Community Ecology, Helmholtz Centre for Environmental Research – UFZ, Theodor‐Lieser‐Straße 4, 06120 Halle, Germany

^4^ German Centre for Integrative Biodiversity Research (iDiv) Halle-Jena-Leipzig, Puschstraße 4, 04103 Leipzig, Germany

^5^ Botanic Garden Meise, Nieuwelaan 38, 1860 Meise, Belgium

^6^ Conservatoire Botanique National Méditerranéen de Porquerolles, 34 avenue Gambetta, 83400 Hyères, France

^7^ Conservatory and Botanic Garden of the City of Geneva, Chemin de l´Impératrice 1, 1296 Chambésy, Geneva, Switzerland

^8^ Plant Evolutionary Ecology, Faculty of Biological Sciences, Goethe University Frankfurt, Max-von-Laue-Str. 13, 60438 Frankfurt am Main, Germany

**Table S1** Sampled study species with details on plant family, the seedbank, region of origin, year of collection of the past population, the number of sampled individuals in 2018, the number of sown seeds and the germination rates for the past (P) and the current (C) populations

| **Species** | **Family** | **Seedbank/Region** | **Collection year past** | **Number of sampled individuals in 2018** | **Number of sown seeds and germination rate P / C** |
| --- | --- | --- | --- | --- | --- |
| *Ammophila arenaria* | Poaceae | CBNMed/Hyères | 1994 | 16 | 100 (20%) / 100 (20%) |
| *Anthemis maritima* | Asteraceae | CBNMed/Hyères | 1992 | 80 | 100 (25%) / 100 (30%) |
| *Anthyllis barba-jovis* | Fabaceae | CBNMed/Hyères | 1992 | 19 | 100 (60%) / 100 (75%) |
| *Elytrigia juncea* | Poaceae | CBNMed/Hyères | 1994 | 25 | 100 (25%) / 100 (45%) |
| *Euphorbia peplis* | Euphorbiaceae | CBNMed/Hyères | 1998 | 20 | 100 (0%) / 100 (0%) |
| *Matthiola tricuspidata* | Brassicaceae | CBNMed/Hyères | 1994 | 15 | 100 (95%) / 100 (98%) |
| *Medicago marina* | Fabaceae | CBNMed/Hyères | 1992 | 59 | 100 (45%) / 100 (90%) |
| *Pallenis maritima* | Plantaginaceae | CBNMed/Hyères | 1994 | 10 | 100 (28%) / 100 (29%) |
| *Plantago crassifolia* | Plantaginaceae | CBNMed/Hyères | 1994 | 10 | 100 (28%) / 100 (29%) |
| *Plantago subulata* | Plantaginaceae | CBNMed/Hyères | 1997 | 103 | 100 (29%) / 100 (78%) |
| *Pseudorlaya pumila* | Apiaceae | CBNMed/Hyères | 1992 | 26 | 100 (0%) / 100 (0%) |
| *Silene nicaeensis* | Caryophyllaceae | CBNMed/Hyères | 1980 | 19 | 100 (1%) / 100 (3%) |
| *Centaurium erythraea* | Gentianaceae | Meise/Namur | 1992 | 20 | 200 (52%) / 1000 (49%) |
| *Clinopodium vulgare* | Lamiaceae | Meise/Namur | 1992 | 47 | 200 (75%) / 200 (97%) |
| *Dianthus carthusianorum* | Caryophyllaceae | Osnabrück | 1993 | 20 | 100 (26%) / 200 (48%) |
| *Digitalis lutea* | Plantaginaceae | Meise/Namur | 1995 | 20 | 500 (20%) / 500 (30%) |
| *Digitalis purpurea* | Plantaginaceae | Meise/ Liège | 1990 | 20 | 200 (100%) / 500 (20%) |
| *Globularia bisnagarica* | Plantaginaceae | Meise/Namur | 1992 | 12 | 100 (49%) / 50 (54%) |
| *Hypericum montanum* | Hypericaceae | Osnabrück/Osnabrück | 1997 | 20 | 250 (20%) / 100 (100%) |
| *Leontodon hispidus* | Asteraceae | Meise/Namur | 1995 | 20 | 300 (32%) / 300 (75%) |
| *Lithospermum officinale* | Boraginaceae | Osnabrück/Osnabrück | 1996 | 20 | 150 (2%) / 150 (2%) |
| *Melica ciliata* | Poaceae | Meise/Namur | 1992 | 21 | 200 (75%) / 150 (50%) |
| *Pimpinella saxifraga* | Apiaceae | Meise/Namur | 1992 | 20 | 200 (50%) / 200 (25%) |
| *Rhinanthus minor* | Orobanchaceae | Meise/ Liège | 1990 | 20 | 200 (1%) / 200 (0.5%) |
| *Sanguisorba minor* | Rosaceae | Meise/Namur | 1992 | 20 | 100 (53%) / 50 (60%) |
| *Sedum album* | Crassulaceae | Meise/Namur | 1992 | 20 | 500 (20%) / 500 (20%) |
| *Silene chlorantha* | Caryophyllaceae | Berlin/Berlin | 1980 | 25 | 150 (32%) / 500 (47%) |
| *Teucrium chamaedrys* | Lamiaceae | Meise/Namur | 1992 | 20 | 200 (20%) / 300 (20%) |

**Table S2** Storage conditions applied by the seed banks

| **Locality** | **Storage conditions** |
| --- | --- |
| Mediterranean species | The seeds of *Plantago subulata* were ultra-desiccated and stored at 17°C, whereas the remaining species were dried and stored at 5°C at CBNMed until we received the seed material in November 2018. |
| Temperate Species | All seeds had been dried at 15% relative humidity and then stored at -20°C at Meise Botanic Garden, the Botanical Garden of the University of Osnabrück and Berlin Botanic Garden and Botanical Museum until we received the seed material in November 2018. |

**Table S3** Results from the statistical cross-species models. (a) Results of linear mixed-effects models testing for the effects of initial size, the year of flowering (flowered in the first year vs. flowered in the second year), the temporal origin (past vs. current population), the region (Mediterranean vs. temperate) and their interactions on the initial size and the standardised flowering onset. (b) Results of generalised mixed-effects models with a binominal distribution testing for the effects of the aforementioned factors on the proportion of flowering individuals and (c) Results of linear models testing for the effects of the 6-year IDM and the 6-year IDM_Diff_ (6-year-De Martonne aridity index), the year of flowering, the temporal origin, the region and their interactions on the standardised flowering onset and the difference in the flowering onset of the two temporal origins, respectively. Significant results (*P*<0.05) are written in bold.

|  | **(a)** | | **(b)** | **(c)** | |
| --- | --- | --- | --- | --- | --- |
|  | **Initial size** | **Standardised flowering onset** | **Proportion of flowering individuals** | **Standardised flowering onset** | **Standardised flowering onset_Diff_** |
| **6- year IDM or  6- year IDM_Diff_** | - | - | - | **F_1_= 7.38 *P*** = **0.015** | **F_1_= 10.43 *P*** = **0.012** |
| **Initial Size** | - | F_1,276.7_= 1.11  *P* = 0.292 | Chi^2^_1_=0.10 *P* = 0.757 | - | - |
| **Flowering Year** | F_1,354.5_= 0.00  *P* = 0.972 | **F_1,12.8_= 9.79 *P* = 0.008** | Chi^2^_1_=0.00 *P* = 0.974 | **F_1_= 8.78 *P*** = **0.009** | F_1_= 0.81 *P* = 0.394 |
| **Temporal Origin** | **F_1,13.7_= 4.45**  ***P* = 0.049** | **F_1,13.8_= 5.60**  ***P* = 0.03** | Chi^2^_1_=1.58 *P* = 0.208 | **F_1_= 8.81 *P*** = **0.009** | - |
| **Region** | F_1,354.41_= 0.00 *P* = 0.948 | F_1,12.8_= 1.81  *P* = 0.202 | Chi^2^_1_=1.79 *P* = 0.181 | F_1_= 0.11 *P* = 0.740 | **F_1_= 6.72 *P* = 0.03** |
| **Initial Size x Lifeform** | - | **F_1,276.7_= 9.34**  ***P* = 0.002** | Chi^2^_1_=0.00 *P* = 0.953 | - | - |
| **Flowering Year x Temporal Origin** | **F_1,13.7_= 10.13**  ***P* = 0.006** | F_1,13.8_= 0.15 *P* = 0.707 | **Chi^2^_1_=5.12**  ***P* = 0.024** | F_1_= 0.38 *P* = 0.544 | - |
| **Flowering Year x Region** | F_1,354.5_= 0.00 *P* = 0.999 | F_1,12.8_= 2.00 *P* = 0.181 | Chi^2^_1_=2.35 *P* = 0.126 | F_1_= 1.25 *P* = 0.279 | F_1_= 0.02 *P* = 0.900 |
| **Temporal Origin x Region** | F_1,13.7_= 0.45 *P* = 0.516 | F_1,13.3_= 2.83  *P* = 0.116 | Chi^2^_1_=1.12 *P* = 0.290 | F_1_= 0.33 *P* = 0.575 | **-** |
| **Flowering Year x Temporal Origin x Region** | F_1,13.7_= 2.51 *P* = 0.136 | F_1,13.3_= 3.44  *P* = 0.086 | Chi^2^_1_=1.44 *P* = 0.230 | F_1_= 1.05 *P* = 0.321 | **-** |

**Methods S1**

**ddRAD library preparation, SNP genotyping and population genomic analyses**

We collected leaf samples from plants grown in a common garden and freeze-dried them. After DNA extraction using the ‘peqGOLD Plant DNA Mini Kit’ (VWR peqlab, Darmstadt, Germany) and DNA quantitation with Qubit (Thermo Fisher Scientific) we followed the ddRAD protocol by (Peterson et al. 2012) with minor modifications, using 100 ng DNA per sample and *EcoRI* and *MspI* as restriction enzymes to generate 12 ddRAD libraries comprising 516 samples of 22 species the multispecies experiment was started with. Libraries were pooled equimolarly and sequenced (PE, 150bp) on four lanes of an Illumina HiSeq2000, resulting in a total of 7.61*10^8^ sequences. The 13 species reported on here comprised 253 samples, between 6 and 10 per species and generation with a total of 3.82*10^8^ sequences.

We used process_radtags from the Stacks 2.0 pipeline (Rochette, Rivera-Colón and Catchen 2019) to demultiplex reads. Sequence data have been deposited in the European Nucleotide Archive (ENA) at EMBL under accession number PRJEB47887 (<https://www.ebi.ac.uk/ena/browser/view/PRJEB47887>) with individual accession numbers ERS7667629 to ERS7668109. Subsequently, we used *dDocent* 2.7.8 (Puritz, Hollenbeck and Gold 2014) to assemble reads to a de novo reference. We set *Clustering_Similarity%* to 0.88, minimum within individual coverage level to include a read for assembly (K1) to 5, minimum number of individuals a read must be present in to include for assembly (K2) to 6 and default values for other parameters. Although the species have different ploidy levels (7 and 6 species are di- and tetra-ploid, respectively) we assumed diploidy for all species because this allowed an identical data analysis across species. Thus, we identified between 1,163,740 and 1,290,150 raw SNPs across species and filtered these raw SNPs following (O’Leary et al. 2018). First, we used the functions vcfallelicprimitives and vcftools to remove indels, keeping only biallelic SNPs with minimum allele count of 3 (*mac* 3), minimum genotype read depth of 3 (*minDP* 3), minimum mean sequence quality of 30 (*minQ* 30), maximum missingness across individuals of 50% (*max_missing* 0.5), skipping fixed heterozygous SNPs based on the observed number of homo- and heterozygotes (*hardy*) and skipping individuals with >75% missing values (*imiss* > 0.75). Subsequently, using vcffilter, we filtered SNPs according to allele balance, strandedness, mapping quality ratio of the two alleles, and status of properly pairing of alleles using parameter values. Using vcftools, we then filtered SNPs to maximum missingness of 33% (*max_missing* 0.66), minimum minor allele frequency (*maf*) of 0.05, minimum mean read depth (*min-meanDP*) of 20 and maximum mean depth (*max-meanDP*) of 1000. In the end we retained only a single SNP per contig. As an exception, we changed parameter settings for *Elytrigia juncea*, the species with the by far largest genome size (25.97 Gb/2C, Zonneveld 2019), and used K1=K2=3, and min-meanDP=5 to increase the number of retained loci. After import into R, we further filtered SNPs to maximum missingness of 30% using gl.filter.callrate (threshold = 0.70) from the dartR package (Gruber et al. 2017). The final data set consisted of 230 samples, between 11 and 20 (average 17.7) per species and between 3 and 10 (average 8.9) per time point, genotyped at between 391 and 2677 (average 1223) bi-allelic SNP loci.

We assessed pairwise genomic relatedness among samples within the two temporal origins using the kinship estimator *r^ß^* (Goudet et al. 2018), through function beta.coan.SNPs available at <https://datadryad.org/stash/dataset/doi:10.5061/dryad.ds8fk04>, which were applied to individuals from the past and the current population in one analysis. The estimator *r^ß^* is a relative measure of relatedness based on genomic marker data which takes a value of zero for pairs of randomly related individuals, positive values for more closely related and negative values for less closely related than expected at random given allele frequencies of the population. For each species, we tested for significant differences of pairwise relatedness between the two temporal populations using analysis of variance (R-function aov).

In addition, we assessed genomic diversity within the past and the current populations as allelic richness, thus correcting for differences in sample size by rarefaction (Ar, El Mousadik and Petit 1996), with the function allel.rich, and the number of private alleles, i.e. alleles that exclusively occurred in either of the past and the current populations, with the function gl.report.pa, both from the R-package PopGenReport (Adamack and Gruber 2014). In case both temporal origins had different sample size, we randomly selected the same number of samples from both populations.

Finally, we quantified neutral genetic differentiation between the past and the current populations as pairwise *F*_ST_ using the function stamppFst from the R-package StAMPP (Pembleton et al. 2013) and tested for significance by bootstrapping 100 times.
